# Differential utilisation of subcellular skeletal muscle glycogen pools: A comparative analysis between 1 and 15 minutes of maximal exercise

**DOI:** 10.1101/2023.09.28.559993

**Authors:** Camilla Tvede Schytz, Niels Ørtenblad, Kasper Degn Gejl, Joachim Nielsen

## Abstract

Distinct subcellular pools of glycogen particles exist within skeletal muscle fibres, distributed both within and between myofibrils and can be found in proximity to, or at a distance from mitochondria. Their precise localisation may influence their degradation rate and role in muscle function. Here, we investigated how exercise at different intensities (1- and 15-min maximal exercise) with known variations in glycogenolytic rate and relative contribution from anaerobic metabolism affects the utilisation of the distinct pools. Further, we investigated how lowered carbohydrate and energy intake affected glycogen volume densities and the storage of glycogen particles (i.e., localisation, size, and number) and their subsequent utilisation during the exercises. Using a randomized, counterbalanced, cross-over design, participants performed two maximal cycle tests of either 1 (n=10) or 15 min (n=10), conducted following consumption of two distinct diets with either high or lowered carbohydrate and energy contents. Muscle biopsies from *m. vastus lateralis* were obtained before and after the exercises. Intermyofibrillar glycogen was preferentially utilised during the 1-min exercise, whereas intramyofibrillar glycogen was preferentially utilised during the 15-min exercise. The lowered carbohydrate and energy intake decreased the particle size across all subcellular pools and reduced the numerical density in the intramyofibrillar and subsarcolemmal pools, with no effects on the glycogen utilisation during the subsequent exercise. In conclusion, the distinct subcellular glycogen pools were differentially utilised during 1-min and 15-min maximal exercise. Additionally, lowered carbohydrate and energy consumption reduces particle size and numerical density, depending on subcellular localisation.

## Introduction

Glycogen serves as a readily available fuel store in human skeletal muscles, stored as particles mainly in three subcellular localisations: intermyofibrillar, intramyofibrillar and subsarcolemmal (Marchand *et al*., 2002). These pools differ substantially in capacity, with the intermyofibrillar pool constituting approximately 80% of the total glycogen content, while the intramyofibrillar and subsarcolemmal pools make up between ∼5-15% (Marchand *et al*., 2002; Nielsen *et al*., 2011). Despite being the minor pools, intramyofibrillar and subsarcolemmal glycogen are preferentially used during exhaustive work lasting 4 to 150 min (Nielsen *et al*., 2011; Gejl *et al*., 2017; Jensen *et al*., 2020b), and in three different models we have found that intramyofibrillar glycogen is closely associated with measures of fatigue resistance (Nielsen *et al*., 2009; Nielsen *et al*., 2014; Jensen *et al*., 2020b). However, these observations are based on work performed with a predominantly aerobic metabolism. Considering that, the energy yield per glycogen particle significantly diminishes during a shift to anaerobic metabolism (Hargreaves & Spriet, 2018), there exist an unmet need to understand how alterations in the ratio between aerobic and anaerobic metabolism, and herein exercise intensity, affect the utilisation of the different pools of glycogen. Due to possible cytoplasmic diffusion restrictions (Verkman, 2002; Ovadi & Saks, 2004) and based on the topological relationships, where intramyofibrillar glycogen is consistently localised distantly from mitochondria, whereas a large proportion of both intermyofibrillar and subsarcolemmal glycogen particles are situated in close proximity to the mitochondria, a shift from aerobic to anaerobic metabolism may have differential effects on the utilisation of these different pools. Therefore, the first aim of the present study was to investigate the effect of maximal exercise of two different intensities on the utilisation of the three subcellular pools of glycogen.

The volumetric density of skeletal muscle glycogen is a product of particle size and numerical density, which both can be modulated (Marchand *et al*., 2007). A glycogen particle is a spherical structure, organized in concentric layers (tiers), comprising both branched and unbranched chains of glucose where each chain gives rise to two chains in the following tier except for the outermost chains (Gunja-Smith *et al*., 1970; Goldsmith *et al*., 1982; Melendez-Hevia *et al*., 1993). Importantly, because of this structure, the size of a glycogen particle is self-limiting (Madsen & Cori, 1958) with the estimated highest possible packing density achieved at the 12^th^ tier (Goldsmith *et al*., 1982; Melendez *et al*., 1998), resulting in a maximal theoretical diameter of 42 nm (Melendez-Hevia *et al*., 1993). Nonetheless, the average size of glycogen particles in human skeletal muscles is submaximal, averaging approximately 25 nm (Wanson & Drochmans, 1968; Marchand *et al*., 2002; Marchand *et al*., 2007; Nielsen *et al*., 2012; Gejl *et al*., 2017; Jensen *et al*., 2021). In support of a preference for medium-sized particles, the numerical density increases without affecting particle size, when glycogen content is increased by a diet- and exercise-induced supercompensation, (Jensen *et al*., 2021). Conversely, when glycogen content is lowered through a carbohydrate (CHO) restricted diet, the decrease is explained by a reduced numerical density (Jensen *et al*., 2021). A theory is that medium-sized particles are optimal for degradation rates due to enhanced accessibility for the glycogen degradation enzymes, achieved by both reducing steric hindrance and increasing the surface-to-volume ratio, although it compromises storing efficiency (Shearer & Graham, 2004), however, this remains to be investigated experimentally. Thus, one inherent determinant of glycogen storage (i.e., particle size vs. numerical density) may entail a trade-off between storing efficiency and metabolic power. Another inherent determinant could be the spatial distribution of the particles (Shearer & Graham, 2004) as this may be important to provide a local provision of energy needed due to diffusion restrictions within the cell (Verkman, 2002; Ovadi & Saks, 2004). These inherent factors might influence how dietary CHO and energy consumption affect the glycogen storage pattern. In the previous mentioned study by Jensen *et al*. (2021) particle sizes in skeletal muscle fibres are remarkable similar across isocaloric diets differing in CHO content, but a low CHO diet reduces the numerical density in all pools. Therefore, the second aim of the present study was to investigate the effect of lowered carbohydrate and energy intake and subsequent maximal exercise at two different intensities, on glycogen volumetric densities, particle size, numerical density, and localisation within skeletal muscle fibres.

Collectively, the aims of the present study were to investigate the effects of maximal exercise at two different intensities (1- and 15-min) as well as lowered energy and carbohydrate intake on glycogen volume densities, particle size, numerical density, and subcellular localisation within skeletal muscle fibres.

## Methods

### Ethical Approval

The human skeletal muscle biopsies included in the present study are a part of a larger study (Schytz *et al*., 2023). This project was approved by The Regional Committees on Health Research Ethics for Southern Denmark (S-20200157) and the experiments conformed to the standards set by the *Declaration of Helsinki* (except for registration in a database). Before providing their written informed consent, the participants were informed about the study and the potential risks and were made aware that they could withdraw from the project at any time.

### Participants

For inclusion the participants had to meet the following criteria: male, age 18 to 40 years, maximal oxygen uptake (VO_2max_) ≥ 45 mL O_2_·min^-1^·kg^-1^, engaging in regular endurance training at least 2 times per week, non-vegetarian, non-smoker, and otherwise healthy. Twenty recreationally active males meeting these criteria were enrolled in the study and randomly allocated to one of two groups performing either 1-min (n=10) or 15-min (n=10) maximal cycling exercise (Table 1). Of note, of the original 12 participants in the 15-min group (Schytz *et al*., 2023), the biopsies from only 10 participants were imaged and included in the current analyses of transmission electron microscopy (TEM) estimated subcellular glycogen pools.

**Table 1.** Participant characteristics.

| Test | n | Age<br>(years) | Body mass<br>(kg) | Height<br>(m) | VO2max<br>(ml min<sup>-1</sup> kg<sup>-1</sup> BM) | POmax inc.<br>(W) |
| --- | --- | --- | --- | --- | --- | --- |
| 1-min | 10 | 23 (22:26) | 78.4 (77.8:79.4)* | 1.85 (1.81:1.88) | 55.4 (54.1;58.6)* | 311 (293:319) |
| 15-min | 10 | 23 (22:23) | 74.4 (70.7:77.4) | 1.82 (1.80:1.84) | 59.3 (55.6;64.2) | 307 (275:326) |

Values are presented as median (interquartile range). \* different from 15-min (*P*=0.004 and *P*=0.049 for body mass and $\dot{V}O_{2max}$ , respectively).

### Study Overview

The twenty participants were randomized to perform two tests of either 1-min (n=10) or 15-min (n=10) maximal cycling exercise, at the end of two dietary intervention periods. In a randomized, and counterbalanced cross-over design the participants, in each of the 1- and 15-min exercise groups, completed two 5-day intervention periods with different CHO consumption strategies with either a high (H-CHO) or a moderate (M-CHO) CHO content (Fig. 1.1). These two strategies were separated by 10 days with habitual exercise and diet. To evaluate the effect of the CHO, energy, and exercise manipulation on the subcellular pools of glycogen, skeletal muscle biopsies were obtained before and immediately after each of the 1-min and 15-min tests (Fig. 1.1).

Each intervention period started with a rest day (Day -4) including a diet consisting of a moderate amount of CHO (M-CHO) (4 g·kg^-1^ body mass (BM)·day^-1^), which was consumed until completion of a glycogen-depleting exercise the following day (Day -3) (Fig. 1.1). Hereafter, the participants were randomized in a counterbalanced order to either proceed on the M-CHO diet or to receive a diet containing a high amount of CHO (H-CHO) (10 g·kg^-1^ BM·day^-1^) for the subsequent part of the intervention period (i.e., Day -3 (after glycogen-depleting exercise), -2, and -1). The participants rested the next day (Day -2) and the following day (Day -1) they completed a short training session consisting of cycling exercise. On the final day (Day 0) of each intervention period the participants ingested a standardised breakfast and completed the performance test (Fig. 1.1).

Within each participant, all procedures were performed at the same time of day, weekday, ergometers, and conducted by the same test leader and standardized verbal encouragement in both intervention periods. Experiments were carried out in the laboratories at the Department of Sport Science and Clinical Biomechanics, University of Southern Denmark, Odense.

### Diet Compositions and Control of Physical Activity Level

A detailed description of diets and physical activity level has previously been described in detail in Schytz *et al*. (2023). In brief, the diet was strictly controlled in the intervention periods, and two distinct diets, denoted M-CHO and H-CHO were administered. These diets contained identical amounts of protein, fat, and dietary fibres (1.6, 1.0 and 0.4 g·kg^-1^ BM·day^-1^, respectively), but differed with respect to CHO content (M-CHO: 4 g·kg^-1^ BM·day^-1^ vs. H-CHO: 10 g·kg^-1^ BM·day^-1^). This difference in CHO content mediated a lower total energy content in the M-CHO diet than the H-CHO diet (138 kJ vs. 240 kJ·kg^-1^ BM·day^-1^). The standardised breakfast, prior to the maximal exercises on Day 0, was ingested two hours preceding the tests (1.3 mL water, 0.9 g CHO, 0.3 g protein, 0.2 g fat and 0.1 g dietary fibre kg^-1^ BM, respectively). The physical activity level was strictly controlled during the intervention periods, and day-to-day diaries confirmed that the participants adhered to these restrictions.

### Preliminary tests

Preliminary tests were conducted ∼1-2 weeks before the first training day (Day -3). The participants were instructed to refrain from strenuous physical activity, alcohol, as well as tea, and coffee from 6:00 pm the day before. Upon arrival to the laboratory, their height and weight (Tanita MC-780MA, Frederiksberg, Denmark) were determined. Then they performed an incremental test to exhaustion to determine VO_2max_ and maximal power output (PO_max inc._) assessments, followed by familiarization to the glycogen depleting protocol and training session (i.e., on Day -3 and -1, respectively). Finally, the participants completed a familiarization trial of either the 1-min or 15-min performance test depending on their random allocation.

The incremental test was performed using a pre-calibrated electromagnetic cycling ergometer (Schoberer Rad Messtechnik (SRM), Julich, Germany) where settings were individualized, and the participants used similar shoes and settings during the glycogen-depleting exercises, training sessions and tests. The test was conducted in a seated position and commenced with a load of 100 W and increased by 25 W every third minute until voluntary exhaustion. A Quark CPET mixing chamber system (Cardio Pulmonary Exercise Test) (COSMED, Albano Laziale, Italy) was used to analyse expired VO_2_ and VCO_2_ in 10-s intervals during the incremental (and maximal exercise tests). Prior to each test, gas-analysers were calibrated with known O_2_ and CO_2_ concentrations, and the flow meter calibrated manually with a 3 L syringe. Moreover, the laboratory was continuously ventilated to maintain constant temperature and composition of the ambient air. VO_2max_ was defined as the highest 30 s average of three consecutive 10 s VO_2_ measurements. PO_max inc._ was determined as the power output of the last completed step plus 25 W times the fraction completed of the last step before reaching VO_2max_.

### Glycogen-depleting exercise and training session

On Day -3 (Fig. 1.1), the participants performed the glycogen depletion exercise protocol consisting of both ergometer cycling (on a pre-calibrated SRM) and arm cranking. The protocol is described in a companion paper (Schytz et al. 2023). Overall, the participants performed a mix of short-term all-out efforts and prolonged continuous exercises. Hereafter, they were randomized in a counterbalanced order to receive either the M-CHO or H-CHO diet. The training session on Day -1 consisted of a 10-min warm-up at 100 W and 4×2 min at 100% of PO_max inc._ separated by 2 min of rest on an air- and magnetic-braked cycling ergometer (Wattbike Pro, Wattbike Ltd. Nottingham, UK).

### 1 min and 15 min of Maximal Cycling Exercise

At test days (Day 0), the participants performed one of the two maximal cycling exercise tests (1-min or 15-min) on the pre-calibrated SRM cycling ergometer. A pre-warmup consisting of 5 min at 100 W followed by 3×1 min at 100% of PO_max inc._, each separated by 1 min of rest, was conducted 30 min before the maximal exercise. Then a muscle biopsy was extracted before the participants warmed up for another 5 min at 100 W. After a short break, the participants increased the cadence to 60 rpm (at a low power output, i.e., <70 W), and the maximal exercise test was then initiated following a 3-s countdown. Cadence was fixed at 110 rpm and 90 rpm in the 1-min and 15-min maximal cycling exercise, respectively. During the 1-min test, power output was not visible for the participants, time was announced orally every 10^th^ second, and the test ended with a 5-s countdown. Prior to the 15-min test, participants were provided a recommended starting load of 90% of PO_max inc._ and was allowed to monitor power output on an adjacent monitor during the first minute of the test. Hereafter, power output was concealed, and time was announced orally every minute during the first 11 min, every 30 s the following 3 min and every 10^th^ second during the last minute, ending with a 5-s countdown. Verbal encouragement was standardised in both tests (i.e., same phrase and tone of voice, and given at the same time-points). Participants remained seated throughout the test and were instructed to generate the highest possible average power output during the test. Technical issues leading to wrong cadence caused the exclusion of one 1-min maximal cycling test in the H-CHO condition. A muscle biopsy was extracted immediately after the performance test. Average power output during the tests and perceived exertion rated immediately after the tests has been reported previously in Schytz *et al*. (2023). Moreover, VO_2_ was measured continuously during the test and used to estimate anaerobic energy contribution (see details below).

### Determination of Anaerobic Contribution and Glycogen Utilisation per External Work

The anaerobic energy contribution was calculated based on estimations of the accumulated oxygen deficit (Medbø *et al*., 1988). First, the individual power-VO_2_ relationship was established based on an average of VO_2_ in the last 60 s of each of the 6-12 submaximal steps of the incremental test. This relationship was used to estimate the VO_2_ needed to cover the energy demand during the 1- and 15- min maximal cycling exercise obtained in 10-s intervals by measurements of power output. The accumulated oxygen deficit for each maximal cycling test was then estimated as the sum of the differences between the calculated O_2_ demand and the measured O_2_ consumption in each 10-s interval. The maximal cycling tests and the incremental test were performed under the same conditions with respect to diet and fluid intake to ensure that differences in the measured VO_2_ was not affected by dietary differences.

The anaerobic energy contribution during the 1- and 15-min maximal cycling exercise was estimated to be on average 64.6% and 4.5% of total energy turnover, respectively (Schytz *et al*., 2023) (NB: the values are re-calculated based on the lower sample size in the 15-min test of the present study). Knowing the relative contributions of energy from aerobic and anaerobic sources enabled a calculation of the ATP yield per glycosyl unit during the 1- and 15-min test assuming a yield of 3 and 36 ATP per glycosyl unit during anaerobic and aerobic conditions, respectively (Hargreaves & Spriet, 2018): 1 min: (36×0.354 + 3×0.646) = 14.7 ATP per glycosyl unit, and 15 min: (36×0.955+3×0.045) = 34.5 ATP per glycosyl unit giving a ratio (1-min/15-min) of 0.43. Thus, due to the larger aerobic contribution during the 15-min exercise and consequently a higher ATP yield per glycosyl unit, a 0.43 times lower amount of glycogen utilisation per external work is expected during the 15-min exercise than during the 1-min exercise. The actual utilisation of the three pools of glycogen (see method below) per amount of external work was calculated as the pre minus the post glycogen value and then divided by the external work performed (i.e., average watts per kg BM times the duration (s)). The values within each maximal exercise for each subcellular region were averaged, and to compare the expected and actual utilisation per external work during the 15-min exercise, the averaged values from the 1-min exercise was multiplied by 0.43, which is indicated by the dotted lines in Fig. 3. This helps determine whether the glycogen utilisation rate is sensitive to the differential contributions from the aerobic and anaerobic metabolism for the specific glycogen pools.

### Muscle Biopsy Extraction and Handling

Muscle biopsies were extracted from the *m. vastus lateralis* before and immediately after (1-2 min) each performance test using the Bergström needle technique with suction through incisions made with scalpels under local anaesthesia (2% lidocaine). The incision for the biopsy extracted immediately after the test was made prior to the test. For each exercise test the pre and post biopsies were extracted from the same leg, changing to the contralateral leg in the next test (i.e., H-CHO or M-CHO condition), with the leg randomly chosen and left, and right leg equally distributed for each diet. The extracted muscle tissue was placed on filter paper on an ice-cooled petri dish, blotted dry and visible connective tissue and fat removed. The muscle tissue was divided in three specimens used in this study: The first specimen was snap frozen in liquid nitrogen and stored at -80°C until biochemical determination of glycogen content. The second specimen was homogenized, frozen in liquid nitrogen, and stored at -80°C for determination of myosin heavy chain (MHC) composition. The third specimen (∼1 mm^3^) was immediately fixed with 2.5% glutaraldehyde in 0.1 M sodium cacodylate buffer (pH 7.3) for 24 h at 5°C, then rinsed 4 x 15 min in 0.1 M sodium cacodylate buffer and stored in 0.1 M sodium cacodylate buffer at 5°C until further processing for TEM. Two post-test biopsies in the 15-min group were not analysed due to technical errors in the tissue preparation phase.

### Biochemical Determination of Glycogen

Muscle glycogen content was determined spectrophotometrically (Beckman DU 650, Beckman Instruments, Fullerton, CA, USA) as previously described in detail in Gejl *et al*. (2014).

### MHC composition

MHC composition was determined as previously described in Schytz *et al*. (2023), and was in brief, determined in homogenate (made with Potter-Elvehjem glass-glass homogenizer (Kontes Glass Industry, Vineland, NJ, USA) from two biopsies (left and right leg) using sodium dodecyl sulfate polyacrylamide gel electrophoresis (SDS-PAGE).

### Preparation of Muscle Specimen for Transmission Electron Microscopy

Muscle specimens were prepared as described by Jensen *et al*. (2022). In brief, muscle specimens were post-fixed with 1% osmium tetroxide and 1.5% potassium ferrocyanide in 0.1 M sodium cacodylate buffer for 120 min at 4°C. Afterwards the muscle specimens were rinsed two times in 0.1 M sodium cacodylate buffer at room temperature, dehydrated through graded series of alcohol at room temperature, infiltrated with graded mixtures of propylene oxide and Epon at room temperature, and embedded in 100% fresh Epon in molds and polymerized at 60°C for 48 h. The blocks of embedded fibres were cut in sections of longitudinally oriented fibres using a Ultracut UCT ultramicrotome (Leica Microsystems, Wetzlar, Germany) in ultra-thin sections (∼60 nm). These sections were contrasted with uranyl acetate and lead citrate, and later examined by TEM.

### Imaging of Muscle Fibers by Transmission Electron Microscopy

The sections were photographed by the same blinded investigator in a pre-calibrated EM 208 transmission electron microscope (Philips, Eindhoven, The Netherlands) with a Megaview III FW camera (Olympus Soft Imaging Solutions, Münster, Germany). From each section (biopsy) 8-10 longitudinal oriented fibres were imaged at 13500× magnification in a randomized but systematic and uniform order (Fig 1.2). From each fibre, 24 images were obtained, including 12 images from the subsarcolemmal region and 12 images from the myofibrillar region hereof 6 images from both the superficial and central parts (Fig 1.2).

### Z-disc-based single fibre analyses

To investigate associations between fibre types and utilisation of the subcellular glycogen pools, the average Z-disc thickness of each fibre was estimated based on 12 random locations (one per myofibrillar image). To ensure that both fibre types are represented, the three fibres demonstrating the thickest and thinnest Z-disc, respectively, were included, leading to an exclusion of 2-4 fibres from each biopsy with intermediate Z-disc widths. The rationale for using Z-disc widths as indicator for fibre type is based on work showing that myofibrillar ATPase properties correlate with Z-disc widths (Sjöström et al. 1982a; 1982b). The analysis was accomplished by the same blinded investigator.

### Quantification of Subcellular Glycogen Volume Densities

Glycogen was quantified in three subcellular localisations within the muscle fibre: 1) the intermyofibrillar space, 2) the intramyofibrillar space, and 3) the subsarcolemmal space (Fig. 1.3) (Jensen *et al*., 2022). The glycogen volume fraction (*V_V_*), taking the effect of section thickness into account, was estimated as proposed by Weibel (1980), assuming that glycogen particles are spherical (Melendez-Hevia *et al*., 1993): *V_V_*=*A_A_* – *t* ((1/π)*B_A_* – *N_A_* [(*t* ·*H*)/(*t* + *H*)]), where *A_A_* is glycogen area density (see estimation below), *t* is the section thickness (i.e., 60 nm), and *H* is the average glycogen particle profile diameter assessed by direct measurement (see below). *N_A_* is the number of particles per area determined by dividing the estimated area density (*A_A_*) by the average glycogen particle area calculated by assuming a circular profile of the glycogen particles and employing the measured mean particle diameter. The glycogen boundary length density is given by: *B_A_*=(π/4)·*S_V_*+*t*·*N_V_*·π·*H*. Here, the surface area density (*S_V_*) can be estimated as: *S_V_*=*N_V_*·*s* where *s* is the mean particle surface area estimated by assuming a spherical particle and the numerical volume density (*N_V_*) is given by, *N_V_*=*N_A_*/(*t*·*H*) (Weibel, 1980).

The glycogen area fraction was estimated by point counting since *A_A_*=*P_P_*, where *P_P_* is the glycogen point density (Weibel, 1980). The intermyofibrillar glycogen content was expressed relative to the myofibrillar space, which mainly consists of the myofibrils, mitochondria, sarcoplasmic reticulum (SR), t-system, glycogen, and lipids. The intramyofibrillar glycogen content was expressed relative to the intramyofibrillar space consisting solely of the myofibrils. Glycogen estimates localised in the superficial region of the myofibrillar space were weighted three times higher than those in the central region since this region takes up 75% of the fibre volume when muscle fibres are assumed to be cylindrically shaped.

The subsarcolemmal region is defined as the region between the sarcolemma and the outermost myofibril consisting mainly of mitochondria, nuclei, glycogen, and lipids. To avoid a bias induced by changes in these parameters during the maximal cycling exercise, the subsarcolemmal glycogen was expressed relative to the muscle fibre surface area, as estimated from length measurement of the outermost myofibril running parallel with the fibre surface membrane multiplied by the sections thickness (i.e., 60 nm).

The total glycogen volume per fibre volume is the sum of all three pools, and since the subsarcolemmal glycogen is expressed as volume per myofiber surface area it was converted to volume per fibre volume. This was obtained by dividing the raw data by 20, since assuming cylindrical fibres, the volume beneath a surface area of 1 µm^2^ is 20 µm^3^ as the volume of a cylinder is: V=0.5·r·A, where r is the fibre radius (i.e., 40 µm) and A is the fibre surface area (here 1 µm^2^).

Grids were overlaid each image for point counting, and different sizes were applied based on expected area fractions to balance the workload of the analysis and the coefficient of error. Thus, grid sizes used for point counting were 400×400 nm, 173×173 nm, 63×63 nm for the intramyofibrillar space, intermyofibrillar glycogen, intramyofibrillar glycogen, respectively, in a frame size of 3.4 µm wide and 2.43 µm long. The grid size was 316×316 nm for subsarcolemmal glycogen. The _est_CE estimated as proposed by Howard and Reed (2005) were 0.12, 0.14, and 0.20 for intermyofibrillar, intramyofibrillar, and subsarcolemmal glycogen, respectively. The mean particle diameter was assessed by measuring the diameter of a minimum of 60 particles per fibre per subcellular localisation (Jensen *et al*., 2022). Only particles with distinct diameter (i.e., no overlapping) were included, and to avoid selection bias included particles were randomly chosen. The average particle volume (size) were estimated by assuming a spherical structure (Melendez-Hevia *et al*., 1993) and employing the average particle diameter for each measurement, while numerical particle density were calculated by dividing the estimated glycogen volume densities by the average particle volume.

Glycogen quantification was done by four trained investigators where all fibres from a participant were analysed by the same investigator. Inter-investigator analyses of 24-35 randomly chosen images showed a bias up to 23% and a coefficient of variation of up to 7% evaluated as proposed by Bland and Altman (1986). The raw data were adjusted for the biases. Also, particle diameters were adjusted for biases of up to 17%. All measurements were blinded and performed in ImageJ (ImageJ 1.53e, National Institutes of Health, USA).

TEM-estimated glycogen was validated against biochemical determined glycogen by plotting the MHC-weighted TEM-estimated total glycogen volume density for each time-point and test against the corresponding biochemically determined glycogen content (Fig 1.4). Including all time points for both tests, a substantial concordance was found (R_c_=0.74) (Fig 1.4), and moderate to substantial concordances were found within time-points for each test (R_c_=0.53-0.74).

### Statistics

Linear mixed models were applied in all analyses because of the repeated measures design, multiple fibres from each biopsy and missing values. Thus, descriptive parameters (i.e., age, body mass, height, VO_2max_, and PO_max inc._) were compared between participants completing either the 1- or 15-min of maximal exercise by including the exercise test as a fixed effect (Table 1). Further, to analyse the effect of exercise test (1- or 15-min) on subcellular glycogen content and relative contribution of each pool to total glycogen, time, test, and time x test interaction were included in the model as fixed effects and participant and time as random effects (Fig. 2). How exercise test affected glycogen utilisation per external work was analysed with test as fixed effect (Fig. 3). The effect of lowering dietary carbohydrates and energy content prior to the maximal exercises (Fig. 7) was examined with diet as fixed effect and participant and fibre type as random effects. The effects of each exercise test (Fig. 8-9) on subcellular glycogen content, relative contribution of each pool to total glycogen, glycogen particle volume and numerical density were examined with time, diet, and time x diet interaction as fixed effects and participant and time as random effects. All main effects and interactions were tested by a Wald test and if significant, pairwise comparisons were conducted.

**Figure 1.**
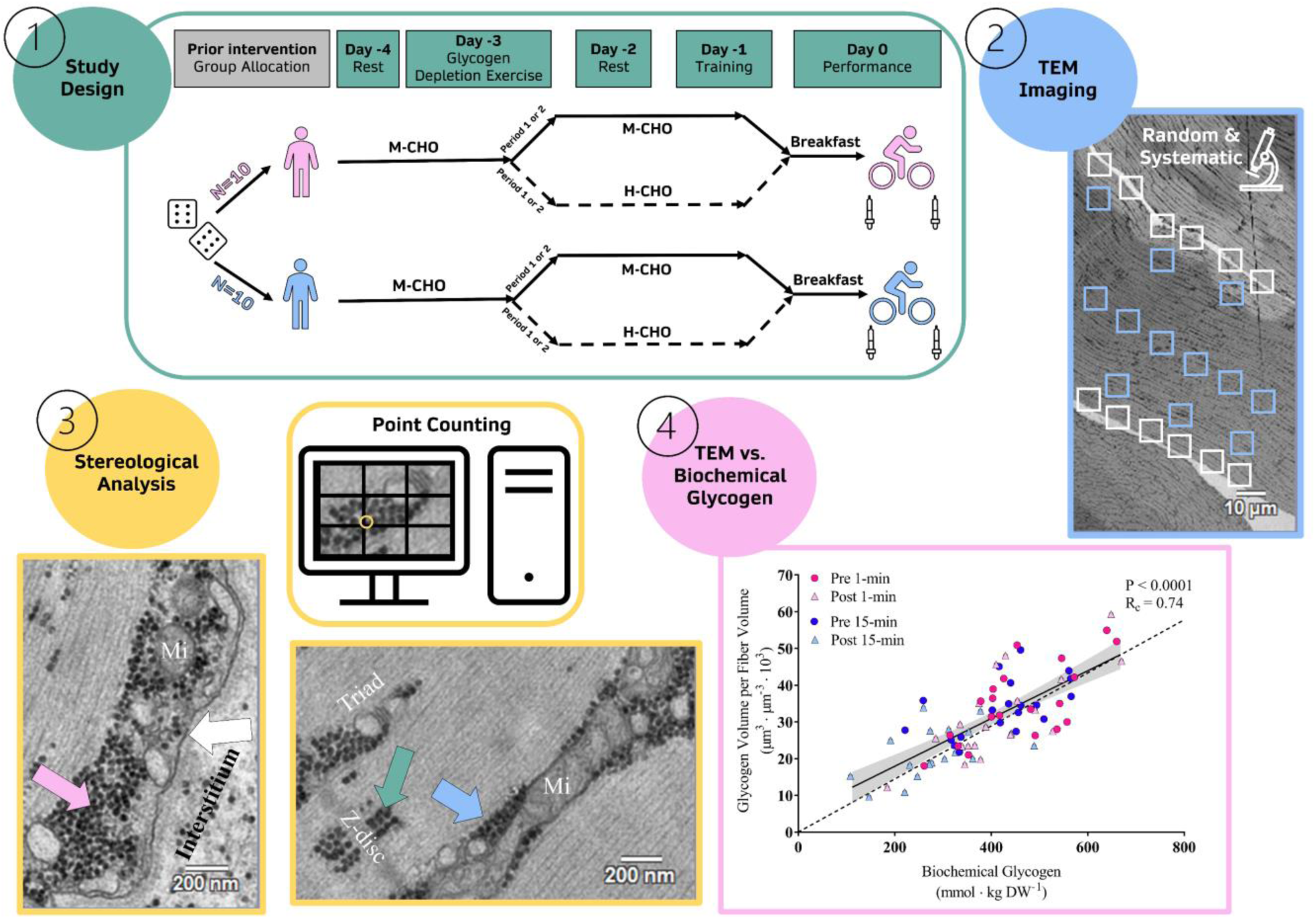
Study Design and Methods. ***1***: The participants were randomised to perform two maximal cycling exercise tests of either 1 min (n=10) (light pink) or 15 min (n=10) (light blue) at the end of two dietary interventions periods consisting of either a moderate (M-CHO) or high (H-CHO) amount of CHO and energy. The participants repeated this after a 10-day period (in a randomised order), where they ingested the opposite diet. Muscle biopsies from the *m. vastus lateralis* were extracted pre and post each exercise test. ***2***: Each longitudinal oriented fibre was photographed by a TEM in a random but systematic order including 12 images from the subsarcolemmal region (white boxes), and 12 images from the myofibrillar region (light blue boxes). ***3***: Glycogen volume density and particle diameter were estimated in the intermyofibrillar (light blue arrow), intramyofibrillar (green arrow), and subsarcolemmal (light pink arrow) localisations. Grids were overlayed each image for manual point counting (example shown with yellow circle). Sarcolemma indicated by white arrow. Example shown of a mitochondrion (Mi), Z-disc, and triad junction consisting of the arrangement of a transverse tubuli with an adjacent terminal cisterna of the sarcoplasmic reticulum on each side. ***4***: TEM-estimated (and MHC-weighted) glycogen content is plotted against biochemical-determined glycogen content. Pre (pink and blue circles) and post (light pink and light blue triangles) values are shown from the maximal cycling exercise of either 1 or 15 min with line of regression and corresponding 95%-CI (grey area). R_c_: Lin’s concordance correlation coefficient based on values normalised to the mean. Dotted line indicating line of identity.

**Figure 2.**
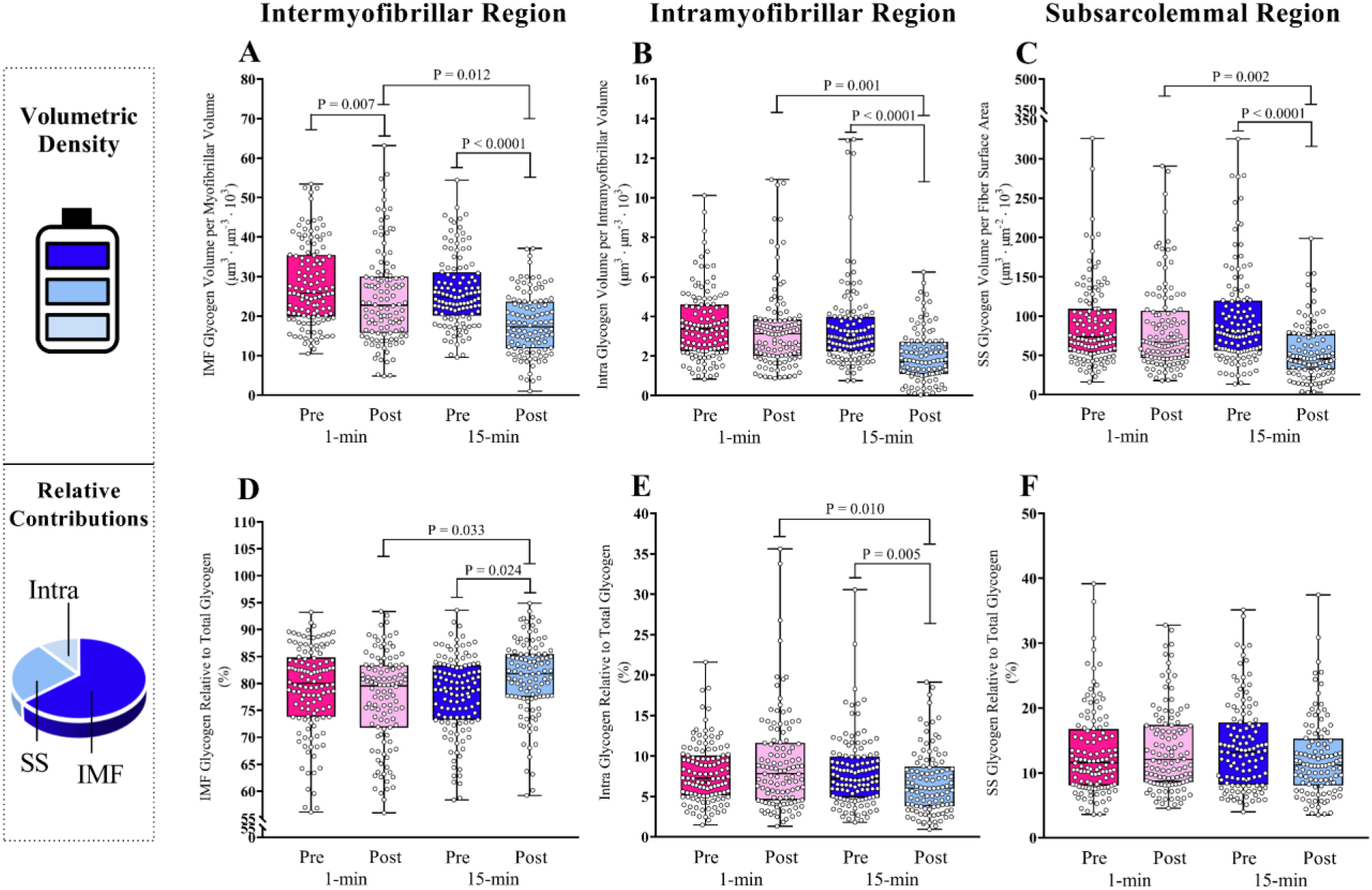
Differential spatial utilisation of muscle glycogen during 1-min versus 15-min maximal cycling exercise. Values are from biopsies obtained before (pre) and immediately after (post) 1-min and 15-min maximal cycling exercise. ***A-C***: glycogen volumetric density; ***D-F***: relative contribution to total glycogen. Intermyofibrillar, intramyofibrillar, and subsarcolemmal abbreviated IMF, Intra, and SS, respectively in the legends. Data shown as median (interquartile range). All individual fibres are displayed as a circle with 6 fibres from each subject. 1-min Pre: n=120, Post: n=114, and 15-min Pre: n=120, Post: n=108. *P*-values represent pairwise comparisons from linear mixed model.

**Figure 3.**
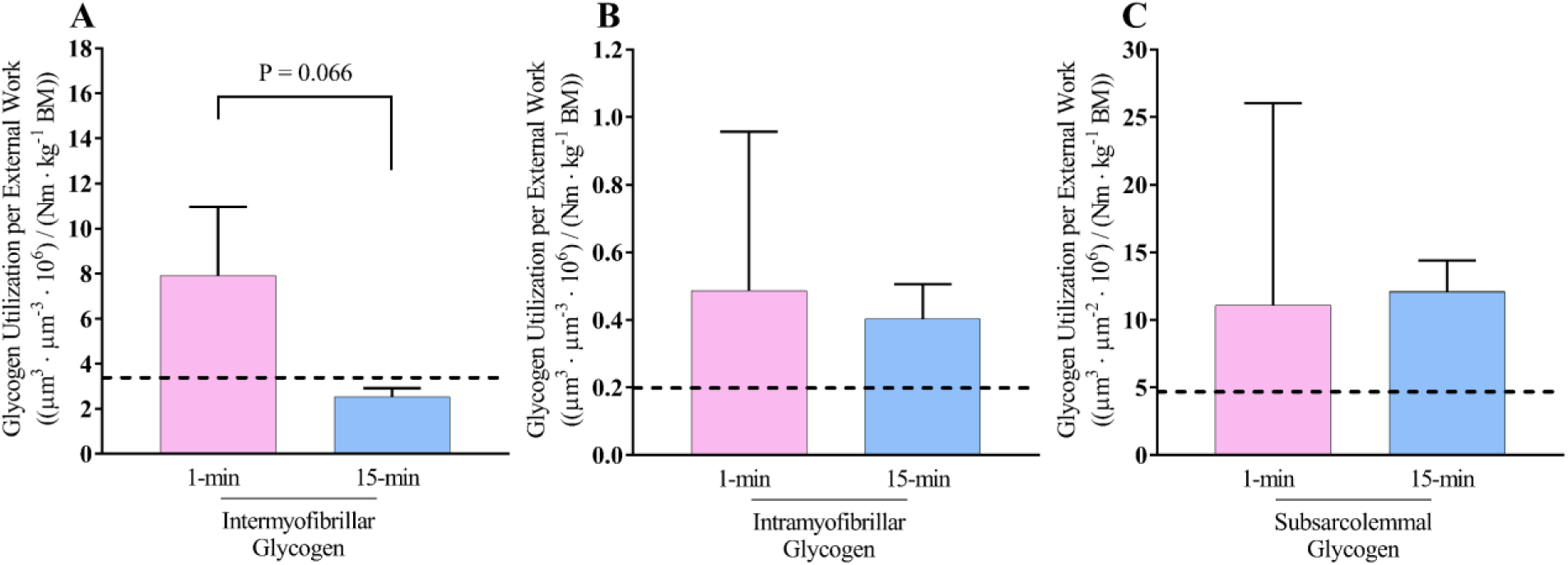
Glycogen utilisation rate per external work in the three subcellular pools of glycogen and during the two different maximal exercise tests. Glycogen utilisation per external work during 1-min (light pink) and 15-min (light blue) maximal cycling exercise in the three subcellular pools: intermyofibrillar (A), intramyofibrillar (B), and subsarcolemmal (C). The black dotted line represents the expected lower glycogen utilisation per external work during the 15-min exercise than during the 1-min exercise if the glycogen utilisation rate is sensitive to the ratio between anaerobic/aerobic metabolism (see Methods for details). Data are presented as mean (standard error of the mean). *P*-value represents the main effect of test from linear mixed model.

Linear mixed models were validated by examining model assumptions: Normal distribution of standardised residuals were investigated by visual inspection of Q-Q plots, and by Shapiro-Wilks’s test and a Skewness and Kurtosis test for normality. Moreover, variance homoscedasticity was investigated by visual inspection of standardised residuals plotted against the predicted values and by running a Breusch–Pagan/Cook–Weisberg test for heteroskedasticity on a linear regression model including all fixed effects for the respective model. If model assumptions were violated data were transformed.

Lin’s concordance correlation coefficient (R_c_) was calculated to validate MHC-weighted total TEM-estimated glycogen against biochemical determined glycogen by testing the agreement with the line of identity, where R_c_ is defined on a scale where 0.21-0.40 is fair, 0.41-0.60 is moderate, 0.61-0.80 is substantial, and 0.81-1.00 is almost perfect concordance (Lin, 1989). This analysis was performed with individual participant values relative to the mean. Correlation analysis between glycogen content in all subcellular regions and Z-disc width were analysed in each exercise test separately and were performed using Pearson’s correlation coefficient. The best fit is stated for each correlation evaluated by the strength of the coefficient of determination (R^2^) evaluated on a scale with <0.20 as weak, 0.21-0.40 as fair, 0.41-0.60 as moderate, 0.61-0.80 as good, and 0.81-1.00 as a very good strength (Fig. 5).

All values are presented as median (interquartile range) unless stated otherwise. Number of participants included in the analysis were 10 in each exercise test. In all statistical tests a significance level of α<0.05 was used. Stata/BE 17.0 (StataCorp, College Station, TX, USA) was applied to conduct the statistical analysis, while figures were constructed in GraphPad Prism 7.05 (GraphPad Software, La Jolla, CA, USA).

## Results

### Utilisation of the subcellular glycogen pools during 1- and 15-min maximal cycling exercise and the relation to the performed external work

Firstly, we investigated across diets if the distinct subcellular pools of glycogen were affected differently by the two exercise tests.

After the 1-min exercise, the largest reduction was observed in the intermyofibrillar pool of glycogen (∼12%), compared to the intramyofibrillar and subsarcolemmal pools of glycogen (∼8-9%; Fig. 2A-C). In contrast, after the 15-min exercise, the largest reductions were observed in the intramyofibrillar and subsarcolemmal pools (∼41-43%), as compared to the intermyofibrillar pool of glycogen (∼30%; Fig 2A-C). Thus, the two exercises had differential effects on the distinct subcellular pools of glycogen. This was also clear if glycogen was expressed as a relative distribution, where the share of intermyofibrillar glycogen increased after the 15-min exercise (time x test: *P*=0.008; Fig. 2D) and intramyofibrillar decreased after the 15-min exercise (time x test: *P*=0.008; Fig. 2E).

The contribution from anaerobic metabolism to the performed external work was estimated (Medbø *et al*. (1988), see Methods) to be 64.6% on average during the 1-min maximal exercise and 4.5% during the 15-min maximal exercise (Schytz *et al*., 2023). With a 12-fold higher ATP yield per glycosyl units by aerobic metabolism than anaerobic metabolism, the glycogen utilisation per external work was expected to be 0.43 times lower during the 15-min exercise than during the 1-min exercise (see Methods). The utilisation of intermyofibrillar glycogen per external work was indeed more than half-fold lower during the 15-min than 1-min maximal cycling exercise, while the utilisation of both intramyofibrillar and subsarcolemmal glycogen per external work was not different between the 1- and 15-min maximal exercises (Fig. 3).

### Single fibre glycogen pools

Fibre-to-fibre variability in the three pools of glycogen (Fig. 4) revealed a higher variability (coefficient of variation) in the volumetric density of intramyofibrillar (0.63) and subsarcolemmal (0.66) glycogen than of intermyofibrillar glycogen (0.43). Single fibre analyses also suggested that the utilisation of glycogen was remarkable homogeneous across fibres in all subcellular pools during 1- and 15-min maximal cycling exercise (Fig. 4). Sorting the single fibres based on the Z-disc width enables a discrimination between fibre types, with type 1 fibres having a relative wider Z-disc than type 2 fibres. Prior to both the 1- and 15-min test a weak negative correlation was found between the Z-disc widths and intermyofibrillar glycogen content suggesting higher pre-exercise glycogen levels in the type 2 fibres, while negligible fibre type differences were observed in the pre-exercise intramyofibrillar and subsarcolemmal glycogen content (Fig. 5A-F). After both 1- and 15-min maximal cycling exercise weak negative correlations were found between the Z-disc width and intermyofibrillar glycogen content, and when comparing the slopes to the post- and pre-exercise correlations a slightly larger utilisation was present in type 2 fibres than type 1 fibres (Fig. 5A and 5D). Also, type 2 fibres may have utilised a larger amount of glycogen than type 1 fibres in the intramyofibrillar region where a weak positive correlation was found after the 1- and 15-min test, while no fibre type differences were observed in the subsarcolemmal region (Fig. 5B-C and 5E-F).

**Figure 4:**
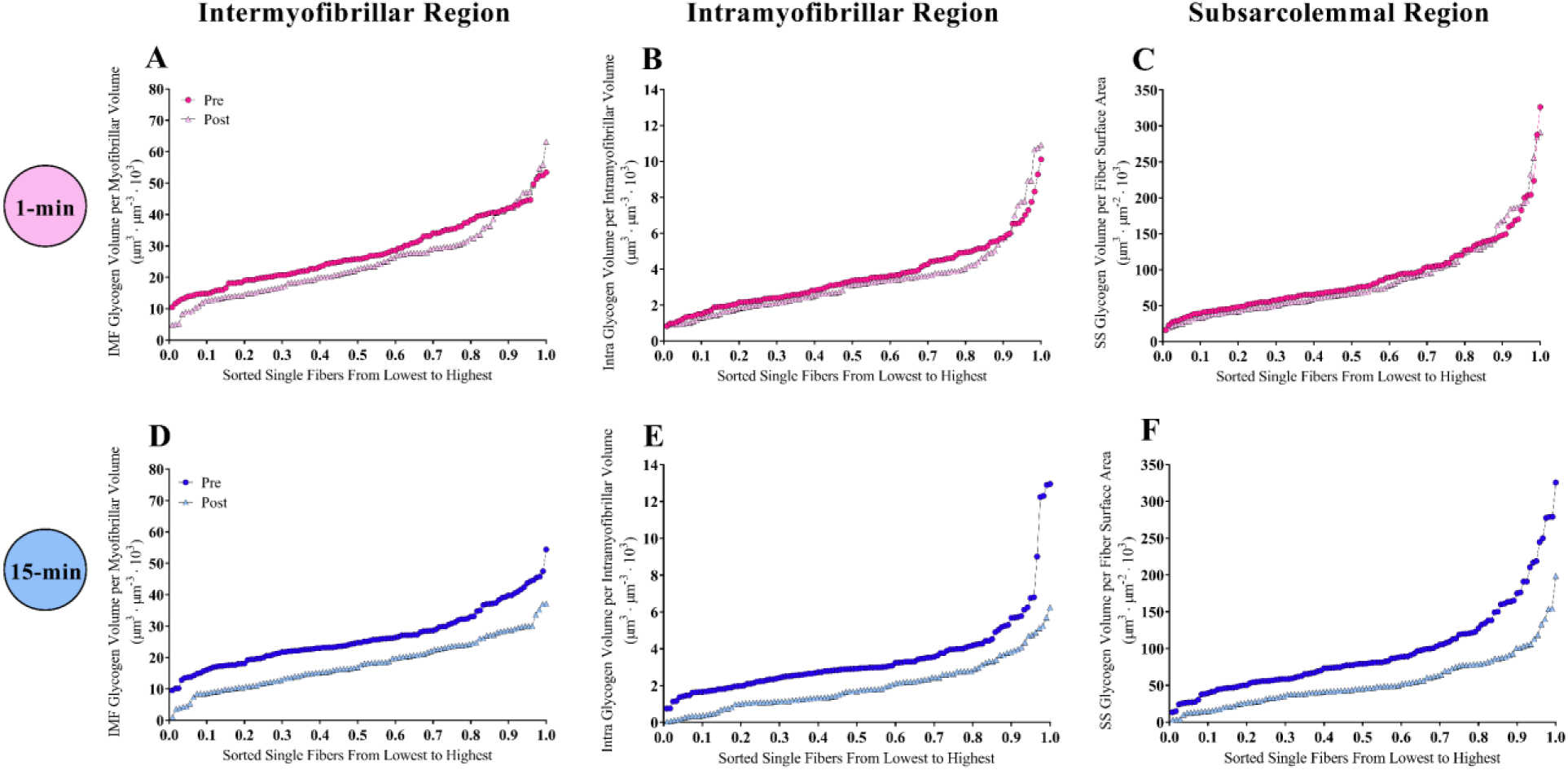
Single fibre glycogen values revealed a fibre-to-fibre homogenous lowering of glycogen during the maximal exercises. Glycogen content of single fibres shown as circles pre (pink and blue) and triangles post (light pink and light blue) 1-min (A-C) and 15-min (D-F) maximal cycling exercise from the intermyofibrillar, intramyofibrillar, and subsarcolemmal region, abbreviated IMF, Intra, and SS in the legends, respectively. Fibers are ordered in relation to glycogen content and weighed according to the number of fibres. 1-min Pre: n=120, Post: n=114, and 15-min Pre: n=120, Post: n=108.

**Figure 5.**
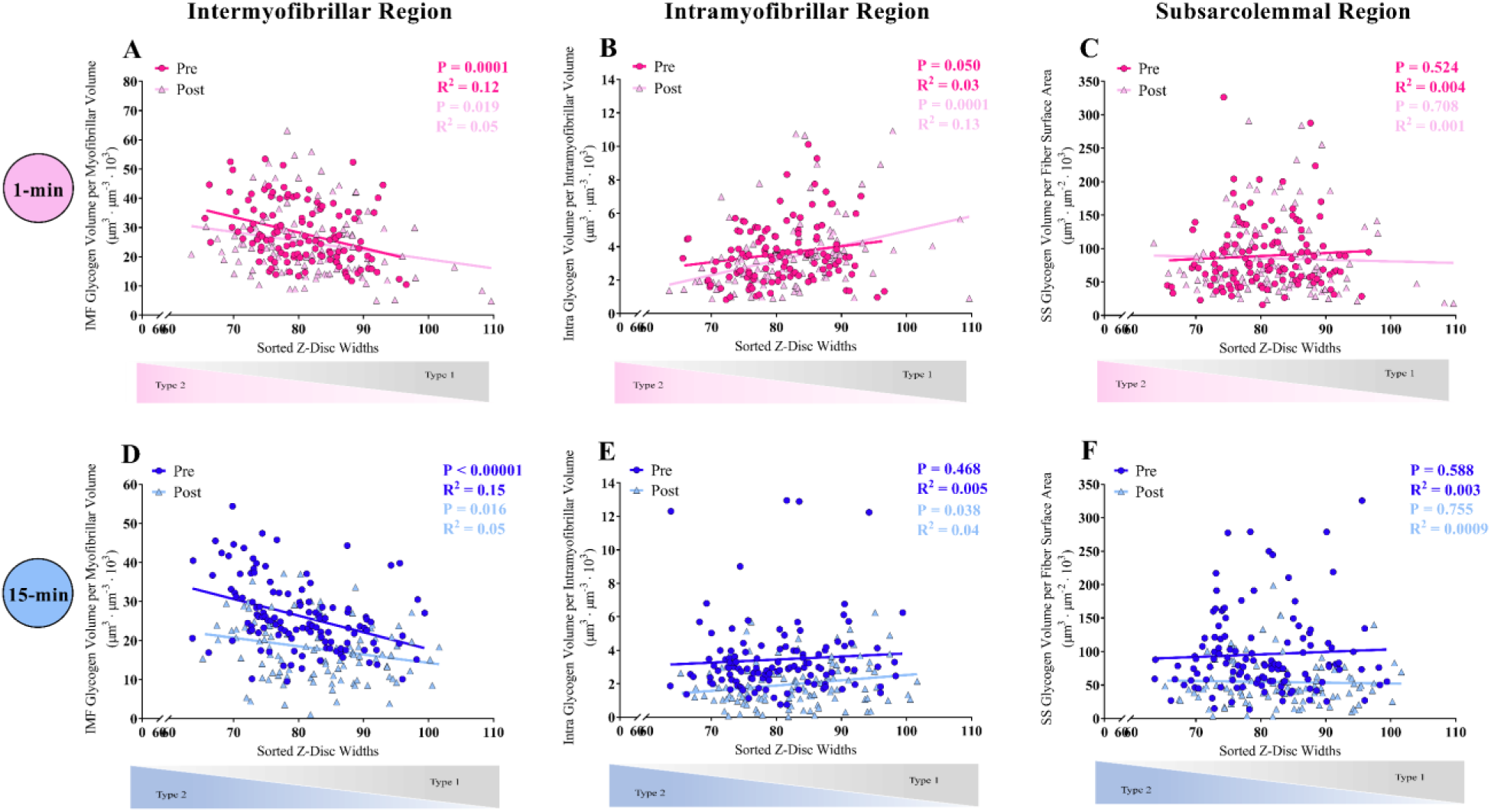
Single fibre Z-disc width suggests a slightly higher glycogen utilisation of type 2 fibres. Glycogen content of single fibres shown as circles pre (pink and blue) and triangles post (light pink and light blue) 1-min (A-C) and 15-min (D-F) maximal cycling exercise from the intermyofibrillar (IMF), intramyofibrillar (Intra), and subsarcolemmal (SS) localisations, respectively. Type 1 fibres have broader Z-disc width than type 2 fibres (See Methods). Fibers are ordered in relation to glycogen content on the y-axis and Z-disc width on the x-axis. 1-min Pre: n=120, Post: n=114, and 15-min Pre: n=120, Post: n=108.

### Effect of lowering carbohydrate and energy intake on pre-exercise glycogen pools, particle size, and numerical density

Lowering the amount of dietary carbohydrates (from 10 to 4 g·kg^-1^BM·day^-1^) equivalent to lowering the total energy intake (from 240 to 138 kJ·kg^-1^BM·day^-1^) in M-CHO, resulted in a 30% lower glycogen content than in H-CHO (367 vs. 525 mmol kg^−1^ DW, respectively) as presented in a companion paper (Schytz *et al*., 2023). The TEM-based analysis of the subcellular glycogen distribution revealed that this diet-induced reduction in glycogen content was most pronounced in the intramyofibrillar and subsarcolemmal pools (Fig. 6A-C), as their relative contribution to total glycogen decreased as opposed to an increase in the contribution of intermyofibrillar glycogen (Fig. 6D-F). The direct measurement of glycogen particle size showed markedly smaller glycogen particles in all subcellular pools in M-CHO compared with H-CHO (Fig. 6G-I), and, to a lesser extent, to fewer glycogen particles in the intramyofibrillar (Fig. 6K) and subsarcolemmal (Fig. 6L) localisations, but not in the intermyofibrillar localisation (Fig. 6J). This unchanged numerical density of intermyofibrillar glycogen particles contributed to a lower diet-induced reduction of glycogen content in this specific localisation, and consequently that intermyofibrillar glycogen constituted a greater part of total glycogen content in M-CHO than H-CHO (Fig. 6D-F).

**Figure 6.**
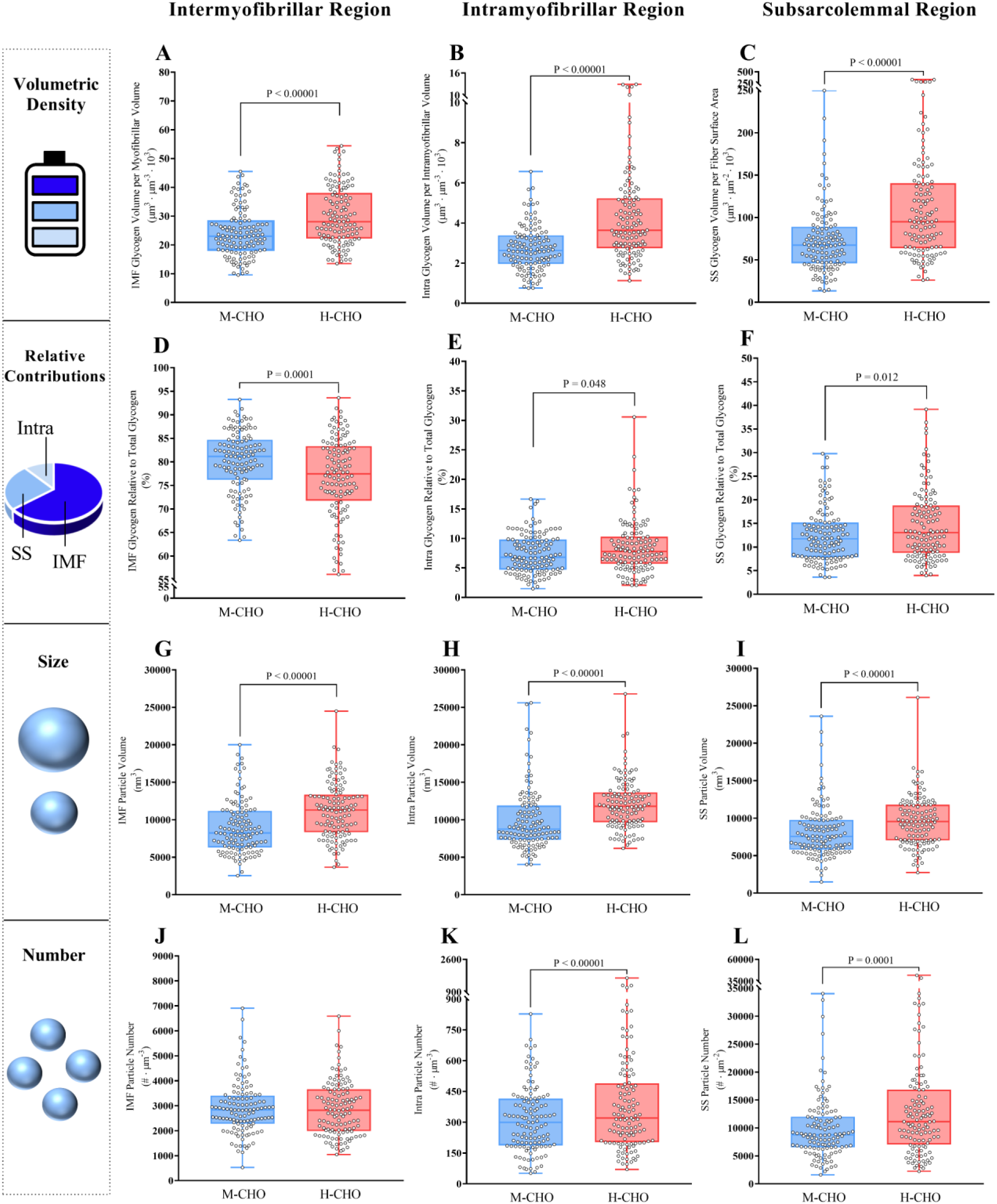
Lowering of the CHO and energy intake reduced the glycogen particle size. All values originate from the biopsies obtained before the exercise. ***A-C***: glycogen volumetric density, ***D-F***: relative contribution to total glycogen, ***G-I***: particle size, ***J-L***: particle number. Intermyofibrillar, intramyofibrillar, and subsarcolemmal abbreviated in the legends as IMF, Intra, and SS, respectively. Data shown as median (interquartile range). All individual fibres are displayed as a circle with six fibres from each participant. M-CHO: n=120 and H-CHO: n=120. *P*-values represent main effects from linear mixed model.

Thus, the lower CHO and energy intake (combined with controlled training sessions) resulted in smaller glycogen particles and localisation-specific fewer particles, with a lower numerical particle density in the intramyofibrillar and subsarcolemmal localisations.

### Effects of 1- or 15-min maximal cycling exercise in the M- or H-CHO conditions on glycogen pools, particle size, and numerical density

We next examined how this altered storage (size, number, and localisation) of glycogen was associated with the utilisation of glycogen during 1-and 15-min maximal cycling exercise. After the 1-min exercise, the volumetric density of intermyofibrillar glycogen was reduced by ∼25% in M- CHO, but unchanged in H-CHO (time x diet: *P*=0.008) (Fig. 7A). Intramyofibrillar and subsarcolemmal glycogen were either unchanged or reduced by up to ∼10-20%, irrespective of diet (time x diet: *P*=0.337 and *P*=0.748, main effect time: *P*=0.137 and *P*=0.219, respectively) (Fig. 7B-C). If the three pools are expressed as a relative distribution of total glycogen content only minor alterations were observed between M-CHO and H-CHO (time x diet: Intermyofibrillar: *P*=0.055, intramyofibrillar: *P*=0.455, and subsarcolemmal: *P*=0.035) (Fig. 7D-F).

**Figure 7:**
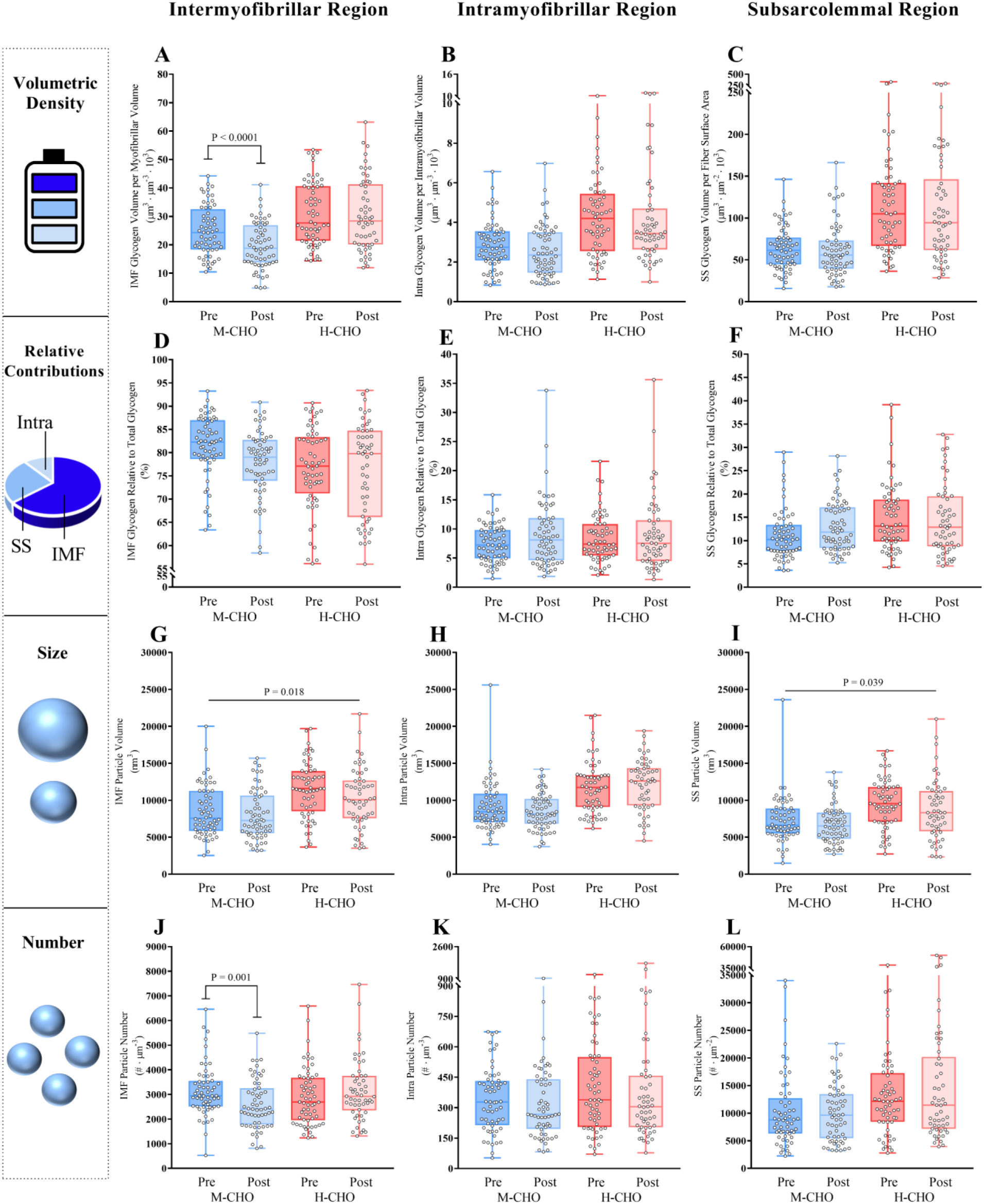
Effects of 1-min maximal cycling exercise on subcellular glycogen utilisation, particle size and number. Values are from biopsies obtained before (pre) and immediately after (post) the 1-min maximal cycling exercise. ***A-C***: glycogen volumetric density, ***D-F***: relative contribution to total glycogen, ***G-I***: particle size, ***J-L***: particle number. Intermyofibrillar, intramyofibrillar, and subsarcolemmal abbreviated in the legends as IMF, Intra, and SS, respectively. Data shown as median (interquartile range). All individual fibres are displayed as a circle with 6 fibres from each subject. M-CHO Pre: n=60, Post: n=60, and H-CHO Pre: n=60, Post: n=54. *P*-values represent main effect of time or pairwise comparisons in case of a time x diet interaction from linear mixed model.

To further understand how the glycogen pools were regulated during 1-min maximal cycling exercise in the M- and H-CHO conditions we investigated the effect on glycogen particle size and numerical density. Interestingly, the reduction in the volumetric density of intermyofibrillar glycogen in M- CHO was primarily driven by a reduction in the numerical density (time x diet: *P*=0.0006) (Fig. 7J), with only a small change in the average particle size (time x diet: *P*=0.505; main effect time: *P*=0.018) (Fig 7G and 9A). The intramyofibrillar and subsarcolemmal glycogen particles did not, or only to a small extent, change in size or numerical density following the 1-min maximal cycling exercise, irrespective of diet (time x diet: intramyofibrillar: *P*=0.108 and 0.854, respectively; subsarcolemmal: *P*=0.465 and 0.878, respectively) (Fig. 7H-I, 7K-L, and 9B-C).

The 15-min maximal cycling exercise mediated reductions in the volumetric density of glycogen in all subcellular pools with no difference between diets (time x diet: *P*=0.272, *P*=0.401, and *P*=0.727, respectively) (Fig. 8A-C). The reductions were most pronounced in the intramyofibrillar and subsarcolemmal localisations than in the intermyofibrillar localisation, resulting in a decrease in the relative contribution of intramyofibrillar glycogen to total glycogen, and an increase in the relative contribution of intermyofibrillar glycogen to total glycogen irrespective of diet (time x diet: *P*=0.240, and *P*=0.786, respectively) (Fig. 8D-E). The glycogen utilisation during 15-min maximal exercise was explained by a decrease in both particle size and numerical density (Fig. 8G-L and 9D-F).

**Figure 8:**
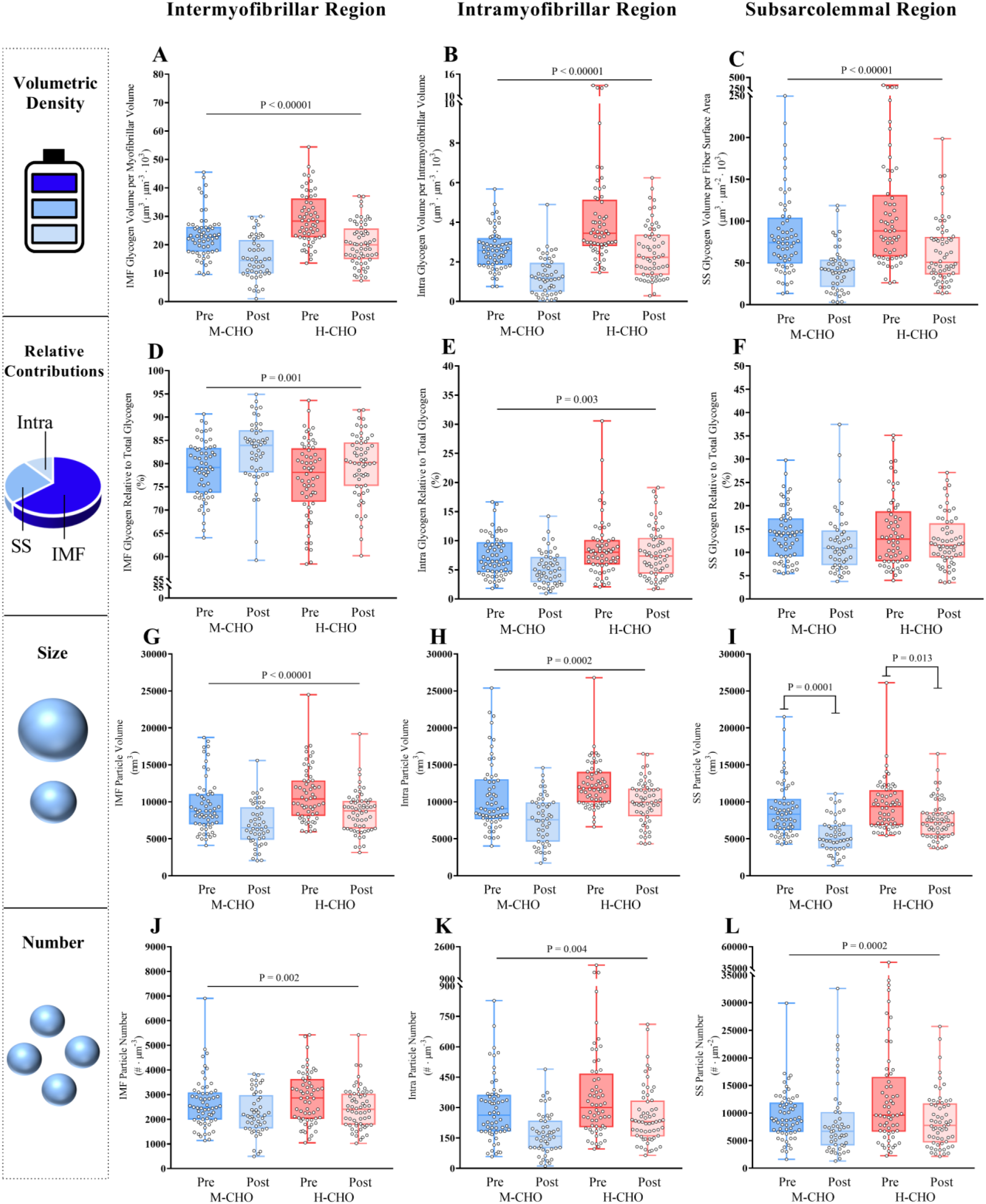
Effects of 15-min maximal cycling exercise on subcellular glycogen utilisation, particle size and number. Values are from biopsies obtained before (pre) and immediately after (post) the 15-min maximal cycling exercise. ***A-C***: glycogen volumetric density, ***D-F***: relative contribution to total glycogen, ***G-I***: particle size, ***J-L***: particle number. Intermyofibrillar, intramyofibrillar, and subsarcolemmal abbreviated in the legends as IMF, Intra, and SS, respectively. Data shown as median (interquartile range). All individual fibres are displayed as a circle with 6 fibres from each subject. M-CHO Pre: n=60, Post: n=48, and H-CHO Pre: n=60, Post: n=60. *P*-values represent main effect of time or pairwise comparisons in case of a time x diet interaction from linear mixed model.

**Figure 9:**
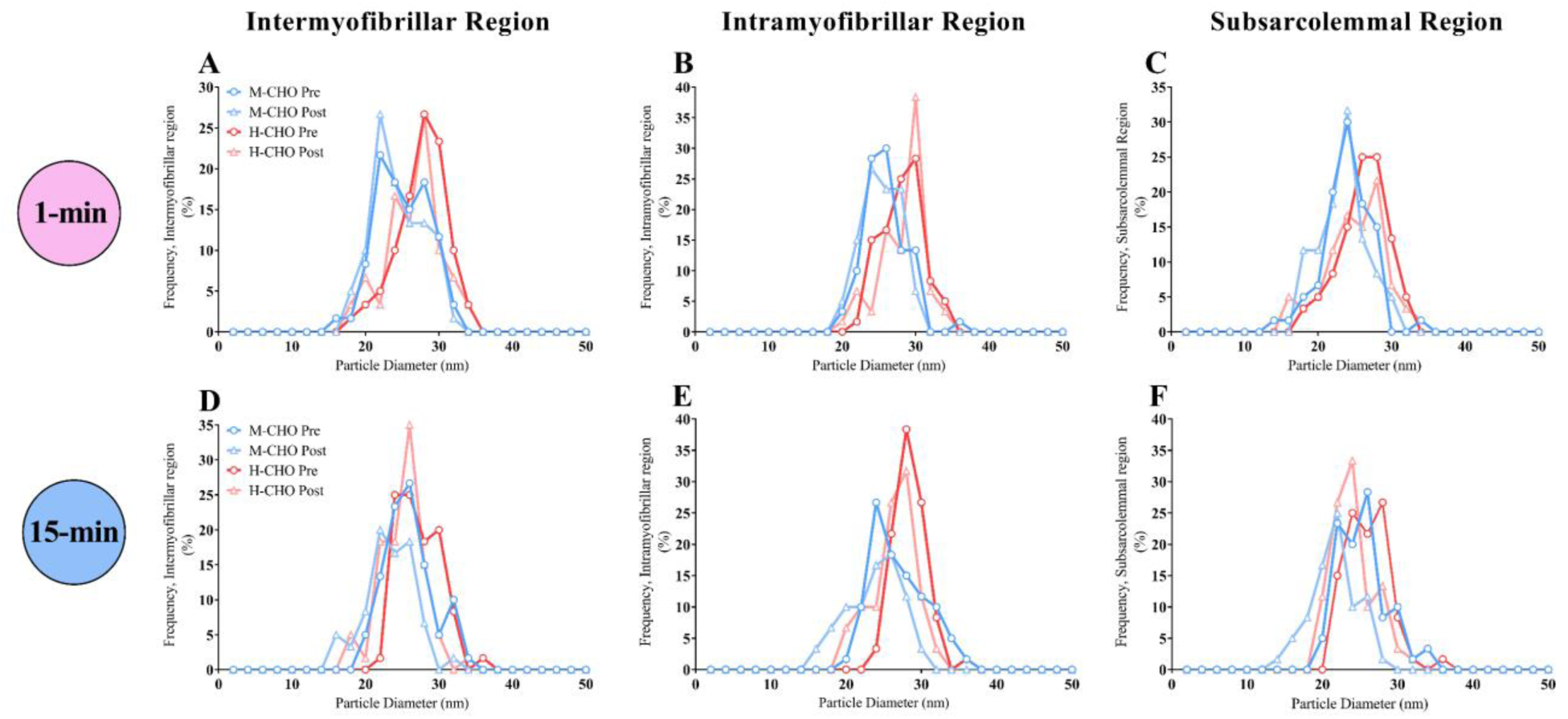
The glycogen particle diameter showed normal distribution. The relative frequency distribution of particle diameter with a bin size of 2 nm in the intermyofibrillar, intramyofibrillar, and subsarcolemmal region prior to and after 1-min (***A***-***C***) and 15-min (***D***-***F***) maximal cycling exercise after adhering to the M-CHO or H-CHO diet. M-CHO Pre (blue circles): n=60/60 (1-min/15-min), Post (light blue triangles): n=60/48, H-CHO Pre (red circles): n=60/60, and Post (light red triangles): n=54/60.

## Discussion

In human skeletal muscle fibres, glycogen is stored as discrete particles distributed heterogeneously throughout the fibre. In this study, we investigated how maximal exercise of two different intensities and lowered carbohydrate and energy intake affected glycogen volume densities, particle size, numerical density, and localisation within skeletal muscle fibres. The main findings of the study were that: 1) the two exercises were characterized by a differential utilisation of subcellular pools of glycogen, where intermyofibrillar glycogen was utilised relatively more than the other pools during 1 min of maximal cycling exercise, as opposed to a relative larger utilisation of intramyofibrillar glycogen during 15 min of maximal cycling exercise; 2) a lowered carbohydrate and energy intake was associated with pronounced smaller glycogen particle sizes in all subcellular pools, and with fewer particles within the intramyofibrillar and subsarcolemmal regions, but not in the intermyofibrillar region; 3) The observed glycogen utilization during the maximal exercises was due to both a decrease in particle size and numerical density, and starting exercise with smaller particles was not associated with any effects on this pattern of glycogen utilisation.

### Differential utilisation of subcellular pools during 1 and 15 min of maximal cycling exercise

During the 1 and 15 min of maximal cycling exercise, differential utilisation of subcellular glycogen was observed, with a relative higher utilisation of intermyofibrillar glycogen during the 1-min exercise and a relative higher utilisation of intramyofibrillar glycogen during the 15-min exercise. The latter observation aligns with previous studies investigating glycogen utilisation during continuous exercise lasting 4 to 150 min (Nielsen *et al*., 2011; Gejl *et al*., 2017; Jensen *et al*., 2020b) or repeated high-intensity intermittent exercise (Vigh-Larsen *et al*., 2022). However, the former finding is, to the best of our knowledge, the first demonstration of this compartmentalised glycogen utilisation pattern in skeletal muscle during maximal exercise of such short duration. Furthermore, analyses of individual skeletal muscle fibres revealed consistently lower glycogen levels across all fibres after compared with before the exercise, and that type 2 fibres (as indicated by the Z-disc width) exhibited a slightly higher utilisation rate than type 1 fibres as expected during such high intensity exercise (Essén, 1978).

The differential utilisation of the three pools of glycogen between the two exercise tests (i.e., intensities and durations) may be attributed to various factors. One of the key distinctions between the two maximal exercise tests is the contribution of anaerobic and aerobic metabolism to the ATP production. Here, we found that during the 1-min exercise, there was a 64.6% contribution from anaerobic metabolism, while in the 15-min exercise, this contribution was only 4.5% (Schytz *et al*., 2023). Given a 12-fold higher yield of ATP per glycosyl unit (glycogen) during aerobic processes than anaerobic processes (Hargreaves & Spriet, 2018), this difference has a substantial impact on the amount of glycogen required.

When comparing the expected and actual glycogen utilisation per external work during the 1 and 15 min of maximal exercise, we observed that the utilisation rates of the intramyofibrillar and subsarcolemmal pool of glycogen per external work performed were not different despite a large difference in the ratio of the energy contribution from aerobic and anaerobic processes. In contrast, the utilisation rate of intermyofibrillar glycogen per external work performed was more than half-fold lower during the 15-min exercise, as expected given the higher aerobic contribution to this type of exercise. Consequently, intermyofibrillar glycogen is preserved during the 15 min of maximal exercise compared to the 1 min of maximal exercise. This preservation is likely due to the increased ATP yield from each glycosyl unit derived from intermyofibrillar glycogen during the aerobic processes. Thus, our findings suggest that during 15-min maximal exercise, metabolites originating from the degradation of intermyofibrillar glycogen can enter the mitochondria and undergo complete combustion. In contrast, metabolites generated through the breakdown of intramyofibrillar and subsarcolemmal glycogen cannot assess the mitochondria. This speculation about compartmentalisation may arise from the proximity between intermyofibrillar glycogen and mitochondria, and the distant localisation of intramyofibrillar glycogen may prevent its interaction with mitochondria. However, subsarcolemmal glycogen particles are also found near mitochondria but showed contrary to the intermyofibrillar glycogen particles no sensitivity to the changes in the aerobic-anaerobic ratio between the 1- and 15-min exercise tests. At present, the role of subsarcolemmal glycogen and mitochondria is unclear. The subsarcolemmal glycogen pool may provide energy for active transport mechanisms across the sarcolemma, which preferentially use glycolytic derived ATP (James *et al*., 1999; Dutka & Lamb, 2007; Jensen *et al*., 2020a). This could explain why this pool of glycogen was insensitive to changes in the anaerobic-aerobic conditions. It has been suggested, however, that energy can be distributed from subsarcolemmal mitochondria to the intermyofibrillar mitochondria through mitochondrial connectivity (Glancy et al. 2015). Thus, subsarcolemmal glycogen could potentially be a source for mitochondrial pyruvate and facilitate energy transfer by intermyofibrillar mitochondria. Nevertheless, our data suggest that this mechanism was not present under the condition of the present project, and it can be speculated that the subsarcolemmal mitochondria were inactive during this type of exercise.

Importantly, calculating the expected glycogen utilisation per external work assumes that all energy provided originate from glycogen. Hence, it is presumed that the contribution of creatine phosphate and blood glucose did not differ between the 1-min and 15-min maximal exercises. The contribution of creatine phosphate is likely 10-20% of the total energy consumption during the 1-min exercise test and negligible during the 15-min exercise test (Gastin, 2001). This discrepancy between the tests will mitigate the anticipated decrease in glycogen utilisation during the 15-minute exercise in comparison to the 1-minute exercise, shifting the estimated value from 0.43 to approximately 0.60. However, this adjustment does not alter the interpretation of the variations in pool-specific glycogen utilisation between the two tests. The assumption about the contribution of blood glucose appears justified, since Katz *et al*. (1986) found almost no contribution of glucose uptake to the metabolism of the working limb during short-term (∼5 min) maximal exercise. This is in contrast to exercise of longer durations where blood glucose would make up a larger part of total energy contribution (Romijn *et al*., 1993), depending on the exogenous supply (Coyle *et al*., 1986). Interestingly, we have previously found that a 4-h recovery period without energy intake predominantly restricts resynthesis of intramyofibrillar glycogen compared to recovery with CHO intake (Nielsen *et al*., 2011), which suggest an association between glucose uptake and intramyofibrillar glycogen. Thus, during exercise of longer durations (>15 min) the glycogen utilisation per external work from the intramyofibrillar pool may decrease due to a larger contribution of blood glucose. In that study subsarcolemmal glycogen was not sensitive to CHO intake, but it can be speculated that the localisation just beneath the sarcolemma makes it sensitive to glucose uptake during prolonged exercise.

### Reducing CHO- and energy-intake mediated smaller pre-exercise glycogen particles

Reducing the amounts of dietary carbohydrates and total energy intake by the M-CHO diet induced a ∼20-30% lower glycogen volume density in the subcellular regions than after ingesting the H-CHO diet. This reduction in glycogen content was primarily mediated by a markedly reduction in glycogen particle size of ∼20-30%, whereas the numerical density of glycogen particles was either maintained or reduced ∼6-21%. Thus, a preservation of the number of glycogen particles seems to be prioritized at the expense of size when storing glycogen under conditions of reduced CHO intake and energy availability.

It is imperative to approach the interpretation of data on particle size and numerical density based on visual inspection of transmission electron micrographs with care, as the smallest particles may not be detectable. As a result, when particles become very small (e.g., due to prolonged exhaustive exercise) it becomes challenging to ascertain any potential decrease in numerical density and the average particle size may be severely overestimated. In the present study, the particles were reduced in apparent size from around 27 nm to 24 nm in diameter. This range in size is well above a potential detection threshold of 8-12 nm, indicating that the number of undetected particles is expected to be minimal.

Nevertheless, the particle diameter was well below the theoretical maximal of 42 nm across all subcellular localisations and after both diets. This is in line with several studies in human skeletal muscles observing that glycogen particles are stored at a submaximal size (Marchand *et al*., 2002; Marchand *et al*., 2007; Nielsen *et al*., 2012; Gejl *et al*., 2017; Jensen *et al*., 2021), possibly to balance the need for a readily available fuel store and efficient storing (Shearer & Graham, 2004). This preference for several medium-sized particles also aligns with an indicated occurrence of smaller glycogen particles, but unaltered total glycogen content, observed when overexpressing glycogenin, the glycogen synthesis-priming protein located in the core of the particle (Skurat *et al*., 1997). Several medium-sized particles also preserves a higher spatial distribution of particles, which could explain the observed down-regulation of glycogen content through the size of particles in the present study.

The difference in particle size between the M-CHO and H-CHO diet contrasts with the observations of Jensen *et al*. (2021). Here, particle sizes were remarkably similar across diets with low, moderate, or high amounts of CHO, while differences in numerical density primarily accounted for differences in glycogen content. This controversy can be interpreted in two ways: 1) there may exist a broad optimum (∼23-27 nm) for the glycogen particle diameter with respect to metabolic power and storing efficiency, since the density of glycosyl units do not vary to a large degree at submaximal sizes as for small and large particles. Accordingly, the numerical density of glycogen particles is not maintained in the low CHO condition in Jensen *et al*. (2021) since the diameter of the particles are close to the lower limit in this optimum (i.e., avoid an unfavourable decrease in particle size with respect to storing efficiency). 2) The diets provided in Jensen *et al*. (2021) were, contrary to the present study, isocaloric, which could indicate that it is the lower energy intake in the M-CHO than the H-CHO diet of the present study that mediated the reduced particle size, rather than the lower CHO-amount *per se*. It remains to be investigated how energy deficiency affects glycogen particle size and numerical density in different subcellular localisations.

A decreased numerical density of intramyofibrillar and subsarcolemmal glycogen particles (but not intermyofibrillar glycogen particles) was observed, leading to a predominantly lowering of the intramyofibrillar and subsarcolemmal glycogen volumetric densities compared with the intermyofibrillar (∼30 vs. ∼20%, respectively). This indicates that these two specific pools of glycogen are more sensitive to dietary carbohydrate intake and/or energy availability than intermyofibrillar glycogen, and that a maintenance of intermyofibrillar glycogen is preferred. A preservation of intermyofibrillar glycogen has also been observed during 4 min of maximal exercise (Gejl *et al*., 2017) and during prolonged exhaustive exercise (Nielsen *et al*., 2011; Jensen *et al*., 2020b). A differential sensitivity of the distinct pools to carbohydrates and/or energy was also found during a 4-h recovery period, with intermyofibrillar glycogen resynthesis favoured in the absence of carbohydrate and/or energy intake, but intramyofibrillar glycogen resynthesis favoured in the presence of carbohydrate and/or energy intake (Nielsen *et al*., 2011). In isolated rodent muscles, we have previously found that intermyofibrillar glycogen correlates inversely with half tetanic relaxation time (Nielsen et al. 2009) and that intermyofibrillar glycogen is the only pool of glycogen, which fuels the SR Ca^2+^ ATPase (Nielsen *et al*., 2022). Thus, several studies suggest a preservation of intermyofibrillar glycogen and this may be a safety mechanism by maintaining a readily energy availability for the SR Ca^2+^ ATPase, and securing re-uptake of Ca^2+^ into SR, and thereby prevent damaging cytosolic Ca^2+^ accumulation (Duncan & Smith, 1978; Turner *et al*., 1988).

### Reducing CHO- and energy-intake did not affect utilisation

Lastly, we investigated how the different storing of glycogen between the H-CHO and M-CHO conditions, in terms of glycogen particle size and numerical density was affected during maximal cycling exercise. In general, it seemed that the pattern of both a decrease in particle size and numerical density was not affected by smaller particles in the M-CHO condition. Previously, we have examined the glycogen particle size before and after 4 min of maximal exercise after ingesting a diet with a high amount of CHO for 24 h (Gejl *et al*., 2017). Here, the average glycogen volumetric density declined more than the glycogen particle volume, indicating that also in this study the utilisation of glycogen was ascribed to a decrease in both particle size and numerical density. We suggest that due to the rather small difference in particle size (∼24 vs ∼27 nm), and, in turn, expected small difference in glycosyl unit density per particle (Melendez *et al*., 1998; Shearer & Graham, 2004) between the M-CHO and H-CHO conditions of the present study, the utilisation pattern (size vs. number) was not affected by differences in pre-exercise particle volume and number.

A surprising finding was that no net reduction of intermyofibrillar glycogen volume density was observed after the 1-min exercise in the H-CHO condition, despite a marked reduction in the M-CHO condition. We interpret this as a less robust finding because biochemically determined glycogen content declined in both conditions, which should be reflected in the intermyofibrillar glycogen volume density, as this pool comprises approximately 80% of the total glycogen content. Thus, the diet intervention is unlikely to have influenced glycogen utilization during the two exercise intensities.

### Concluding remarks

We utilised transmission electron microscopy and standard stereological techniques to quantify the volumetric density of skeletal muscle glycogen within three distinct subcellular pools. Biopsies were extracted from young males subjected to either 1 or 15 minutes of maximal ergometer cycling exercise. Our analysis revealed divergent preferences for glycogen storage pools during the two exercise durations. Specifically, 1-minute maximal exercise predominantly depleted intermyofibrillar glycogen, whereas 15-minute maximal exercise primarily targeted intramyofibrillar glycogen. As exercise duration extends from 1 to 15 minutes, the most conspicuous disparity observed is the heightened contribution of aerobic metabolism to energy provisioning. The findings from this investigation propose that the degradation rate of glycogen ceases exclusively for intermyofibrillar glycogen when transitioning from 1-minute to 15-minute maximal exercise, while the degradation rate of intramyofibrillar and subsarcolemmal glycogen remains elevated. If substantiated, these results indicate that regulation of glycogenolytic rate during exercise is dependent on the subcellular localisation in human skeletal muscle fibres. Also, our analysis revealed that a lowering of dietary carbohydrates and energy induced a lower glycogen particle size across all subcellular pools, while the numerical density was lower in the intramyofibrillar and subsarcolemmal pools. Noteworthy, this disparate storing pattern had no effect on the subsequent utilisation during the maximal exercises.

## Acknowledgments

We thank Dorte Mengers Flindt, Chris Christensen, Mikkel Egeskov Vigen and Martin Harris Larsen for technical assistance. The imaging by transmission electron microscopy was performed at the Core Facility for Integrated Microscopy, Faculty of Health and Medical Science, University of Copenhagen.

## Author Contributions

CTS, KDG, NØ and JN contributed to the conception and design of the experiments. All authors contributed to the data collection, analyses and/or data interpretation. CTS and JN drafted the manuscript, while all authors edited and/or revised the manuscript. The final version of the manuscript was approved by all authors, and all agree to be accountable for all aspects of the work in ensuring that questions related to the accuracy or integrity of any part of the work are appropriately investigated and resolved. All persons stated as authors qualify for authorship, and all those who qualify for authorship are listed.

